# Meta-microRNAs as potential noninvasive markers for early diagnosis of Alzheimer’s disease

**DOI:** 10.1101/281915

**Authors:** Ning Jiang, Can-Jun Ruan, Xiao-Rui Cheng, Lu-Ning Wang, Ji-Ping Tan, Wei-Shan Wang, Fang Liu, Wen-Xia Zhou, Yong-Xiang Zhang

## Abstract

**Introduction:** The aim of this study was to investigate the potential role of a microRNA panel as early diagnostic markers for Alzheimer’s disease (AD).

**Methods:** The differentially expressed serum microRNAs were screened with microarray among cognitively normal controls (CNC), mild cognitive impairment (MCI), and AD. QRT-PCR assay was applied to evaluate differentially expressed microRNAs with two independent cohorts including 202 participants. Logistic regression model based on microRNA panel was constructed using a training cohort and then validated using an independent cohort.

**Results:** First, four differentially expressed serum microRNAs (let-7g, miR-197, miR-126 and miR-29a) were found, which expressions were positively correlated with mini mental state examination (MMSE) score. Second, a microRNA panel with the four microRNAs demonstrated good diagnostic performance for MCI and AD with 84% and 92% accuracy. Third, when combined with MMSE score, the diagnostic performance of the microRNA panel was further improved.

**Discussion:** Blood microRNAs are potential AD biomarkers that may lead to new diagnostic strategies.

## Introduction

Alzheimer’s disease (AD) is an age-related neurodegenerative disorder that is characterized by progressive memory loss and deteriorated higher cognitive functions. Increasing evidence suggests that the pathophysiological process of AD begins well before the diagnosis of clinical dementia (Ritchie et al, 2016). The majority of AD patients are asymptomatic during the preclinical stages of the pathological process which is believed to be a period of approximately 17 years (Villemagne et al, 2013). Therefore, development of early and accurate diagnosis for AD is crucial, enabling accurate diagnoses and facilitating the possibility for an early and specialized treatment to slow or delay the progression of the disease and provide prognostic information. Current methods for diagnosing AD involve a detailed history and neuropsychological testing to establish the presence of dementia, and a definitive diagnosis of AD can only be made postmortem. The new diagnostic methods based on imaging techniques and analysis of proteins and other components in the cerebrospinal fluid (CSF) have been developed recently (Fagan et al, 2011; Sepulcre & Masdeu, 2016; Wang et al, 2015). These methods, however, are not suitable for the everyday clinical screening due to their invasiveness and relatively high cost. Thus, the identification of reliable, non-invasive and inexpensive biomarkers is a major challenge before or during the pre-clinical phase in order to diagnose and monitor AD progression and enable early treatments to slow or delay the progression of the disease.

Mild cognitive impairment (MCI) is defined as the intermediary state between normal old age cognition and AD (Winblad et al, 2004), which reflects preclinical AD in a considerable portion of the affected individuals. Patients with MCI have high risk of developing AD with conversion rates about 12% annually and 80% at six years follow-up (Petersen, 2004). Consequently MCI may be a useful AD prodromal phase in which to test putative biomarkers for their efficacy in early disease detection.

MicroRNAs are a group of small noncoding RNAs that modify gene expression at the post-transcriptional level. Thus, microRNAs are important epigenetic regulators of numerous cellular processes (Bartel, 2009; McNeill & Van Vactor, 2012). In the past years, microRNAs have gained increasing attention in studies on neurodegenerative diseases. Recently, circulating microRNA levels have been proposed as potential diagnostic tools for a number of diseases (Wang et al, 2010; Zhou et al, 2011). Accumulating evidences of microRNA expression profiles reported by several groups showed that microRNAs were involved in AD progressions (Muller et al, 2014; van Harten et al, 2015) and altered in response to Aβ indicating that microRNA and AD had a direct link (Schonrock et al, 2010; Sorensen et al, 2016). However, those studies were limited by one or more of the following factors: limited number of screened microRNAs, small sample size, and lack of independent validation. Importantly, cell-free microRNAs have been shown to be stable in blood samples (Mitchell et al, 2008), which laid the foundation for studying the role of serum/plasma microRNAs in the diagnosis and prognosis of disease.

In this study, we aimed to investigate serum microRNA expression profiles (877 microRNAs) with independent validation in a large cohort of 202 participants, with the intention to identify a panel of microRNAs for the early diagnosis of AD, thus to explore their clinical significance in disease development and progression, and provide information for personalized therapy.

## Results

### MicroRNA screening and testing

The strategy of identifying differential microRNAs for early diagnosis of AD in this study was to choose candidate microRNAs on the basis of pairwise comparison of AD versus cognitive normal control (CNC), AD versus MCI, and MCI versus CNC, respectively. The serum samples that met the eligibility criteria were allocated to the discovery, training and validation phases (Figure 1). The characteristics of study participants in three phases were listed in Table 1. Expression profiling of microRNAs of sixty serum samples in CNC, MCI and AD was performed using the Agilent microRNA microarrays containing 877 human microRNA probe sets, which has been shown to produce precise and accurate measurements of microRNA levels that span a linear dynamic range from 0.2 amol to 2 fmol of input microRNA and good correlation to Taqman assay (Fumiaki Sato, 2009). Hierarchical clustering of microRNA expression profiles in three group comparisons was illustrated in Supplement Figure 1. A Mann-Whitney test was performed to discover differentially expressed microRNAs in the three pairwise comparisons: MCI and AD versus CNC respectively. Significance analysis resulted in the identification of eighty two microRNAs as differentially expressed among the CNC, MCI and AD groups (Supplement Table 1A-1C). From the differentially expressed microRNAs, 29 detectable microRNAs with *P* value < 0.05 and fold expression change > 2 were identified between the MCI and CNC groups (Supplement Table 1A), 23 detectable microRNAs with *P* value < 0.05 and fold expression change > 2 were identified between the AD and CNC groups (Supplement Table 1B), and 30 detectable microRNAs with *P* value < 0.001 and fold expression change > 2 were identified between the AD and MCI groups (Supplement Table 1C). There was twelve microRNAs overlapping between the three group comparisons. Finally, 14 microRNAs with *P* value < 0.037 and fold change > 2.6 from the 70 differentially expressed microRNAs were selected for further validation by quantitative reverse transcriptase polymerase chain reaction (qRT-PCR), with cohort of 30 participants (Supplement Table 1D).

**Table 1:**
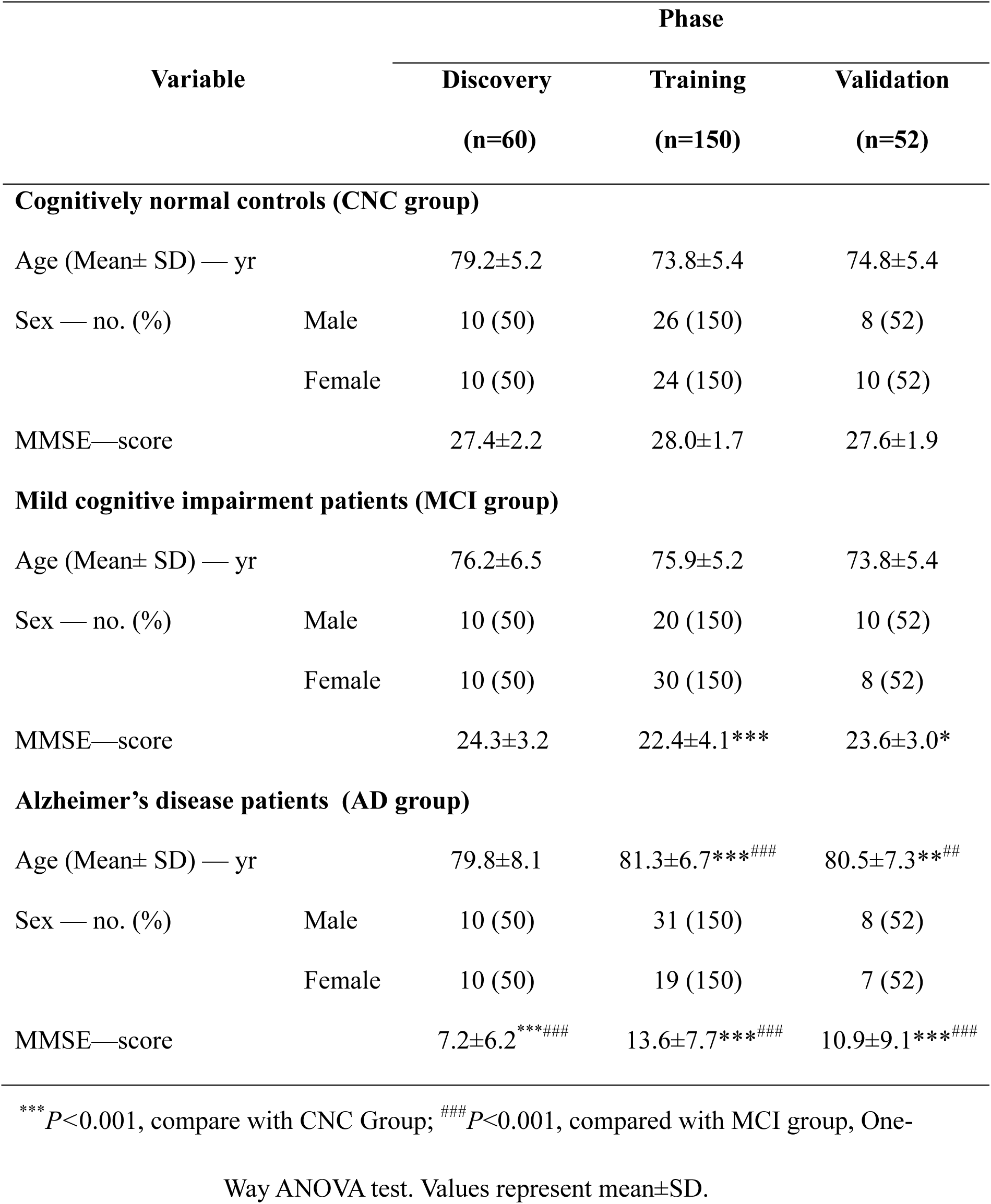
Characteristics of study subjects in the Discovery, training and validation Datasets

**Figure 1:**
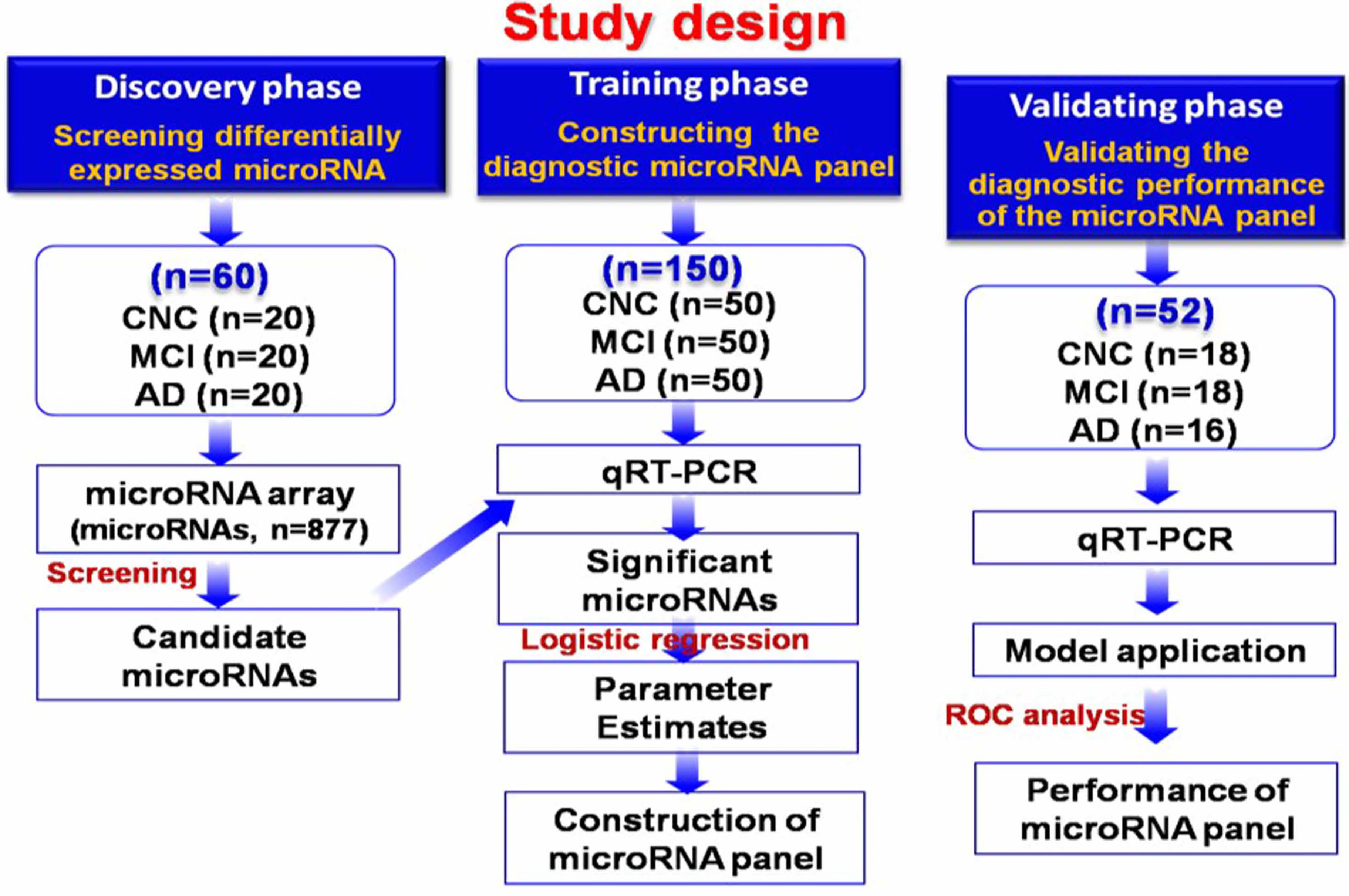
Study design. CNC, cognitive normal control; MCI, mild cognitive impairment; AD, alzheimer’s disease; ROC, receiver operating characteristics; RT-PCR, reverse transcriptase polymerase chain reaction.

### Differential expression profile of four candidate microRNAs

The 14 selected microRNAs were first tested using an independent cohort of 30 serum samples with qRT-PCR, and five microRNAs (let-7g, miR-197, miR-126, miR-29a and miR-451) of 14 passed the quality control. The expression levels of four microRNAs (let-7g, miR-197, miR-126 and miR-29a) were significantly lower in the serum of AD and MCI when compared with CNC group (Figure 2). The expression profile of those four individual microRNAs was further evaluated with qRT-PCR on 120 additional serum samples. The combined 150 serum samples were used as the training data set for the construction of the diagnostic microRNA panel for the differentiation not only between the AD group and the CNC group but also between the MCI group and the CNC group.

**Figure 2:**
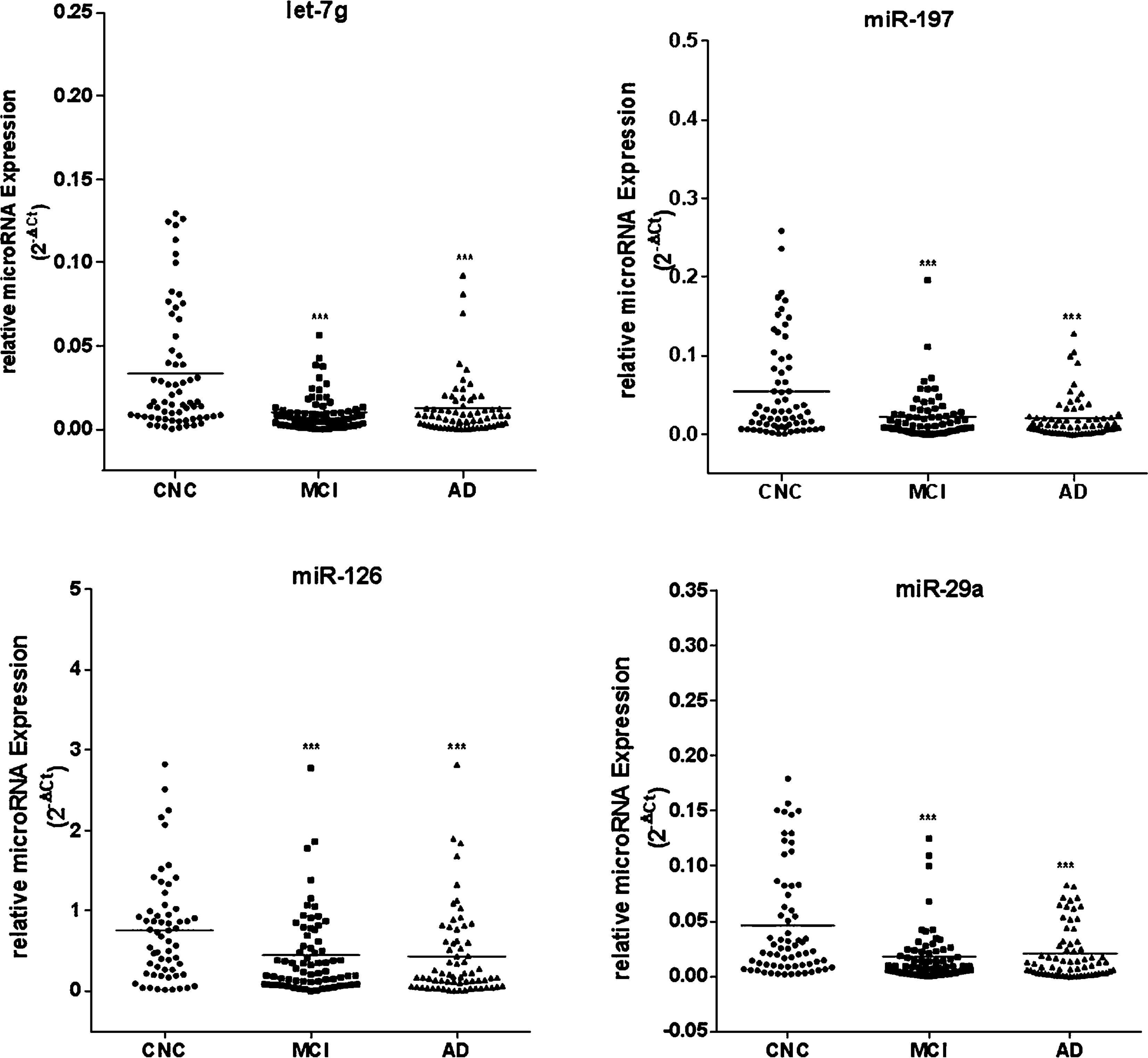
The expression levels of let-7g, miR-197, miR-126 and miR-29a in serum of CNC, MCI and AD patients. The expression levels of serum let-7g, miR-197, miR-126 and miR-29a were significantly lower in MCI and AD compared with CNC (*P*<0.05 ∽0.0001). Values represent mean ± SEM. CNC n=50, MCI n=50, AD n=50. ^***^ *P*<0.0001, ^**^ *P*<0.001, ^*^ *P*<0.05, versus CNC group. One-Way ANOVA with Kruskal-Wallis test.

### MicroRNA expression profile for MCI versus CNC and AD versus CNC in the training data set

The four differentially expressed microRNAs (let-7g, miR-197, miR-126, miR-29a) were evaluated in serum samples of the training data set (150 participants). Scatter-plots of expressions of the four microRNAs in CNC, MCI, and AD were illustrated in Figure 2. Lower expression levels of let-7g, miR-197, miR-126, and miR-29a were observed in patients with MCI compared with those in CNC group (*P*<0.0001 or *P*<0.001) (Figure 2) and fold changes were 0.3, 0.3, 0.4, and 0.3, respectively (Supplement Table 2). The accuracy of let-7g, miR-197, miR-126 and miR-29a for diagnosing MCI from CNC, measured by AUC, was 0.71, 0.68, 0.68, and 0.72, respectively, and the multivariate *P* values for both miR-197 and miR-126 were <0.05 (Supplement Table 2). Similar trend of lower microRNA expressions was also found in patients with AD compared with those in CNC group (*P*<0.0001 ∽0.05) (Figure 2) and fold changes were 0.3, 0.3, 0.4, and 0.4 for let-7g, miR-197, miR-126, and miR-29a, respectively (Supplement Table 3). The corresponding AUCs of let-7g, miR-197, miR-126 and miR-29a in differentiating AD from CNC were 0.71, 0.71, 0.70, and 0.67, respectively (Supplement Table 3). Because age is acting as confounding factor in AD, the relation between age and expression of the four microRNAs was studied with the Pearson correlation analysis. The results showed that there were no significant correlations of the expressions of let-7g, miR-197, miR-126 and miR-29a with age (Figure 3) and Pearson r was -0.13, -0.11, -0.01, and -0.05 (*P*>0.05), which indicated the four microRNAs might have no relations with age. To examine the relation of microRNA expression and cognitive function, the Pearson correlation analysis among the expressions of let-7g, miR-197, miR-126 and miR-29aand MMSE scores were performed. Significant positive correlations of the expressions of let-7g, miR-197, miR-126 and miR-29a with MMSE score were found (Figure 4) and Pearson r was 0.29, 0.25, 0.28 and 0.22 (*P*<0.0005, 0.001, 0.0001 and 0.001, respectively). The results suggested that the four microRNAs might have relations with cognitive function and they turned out to be potential predictors for differentiating MCI and AD from CNC.

**Figure 3:**
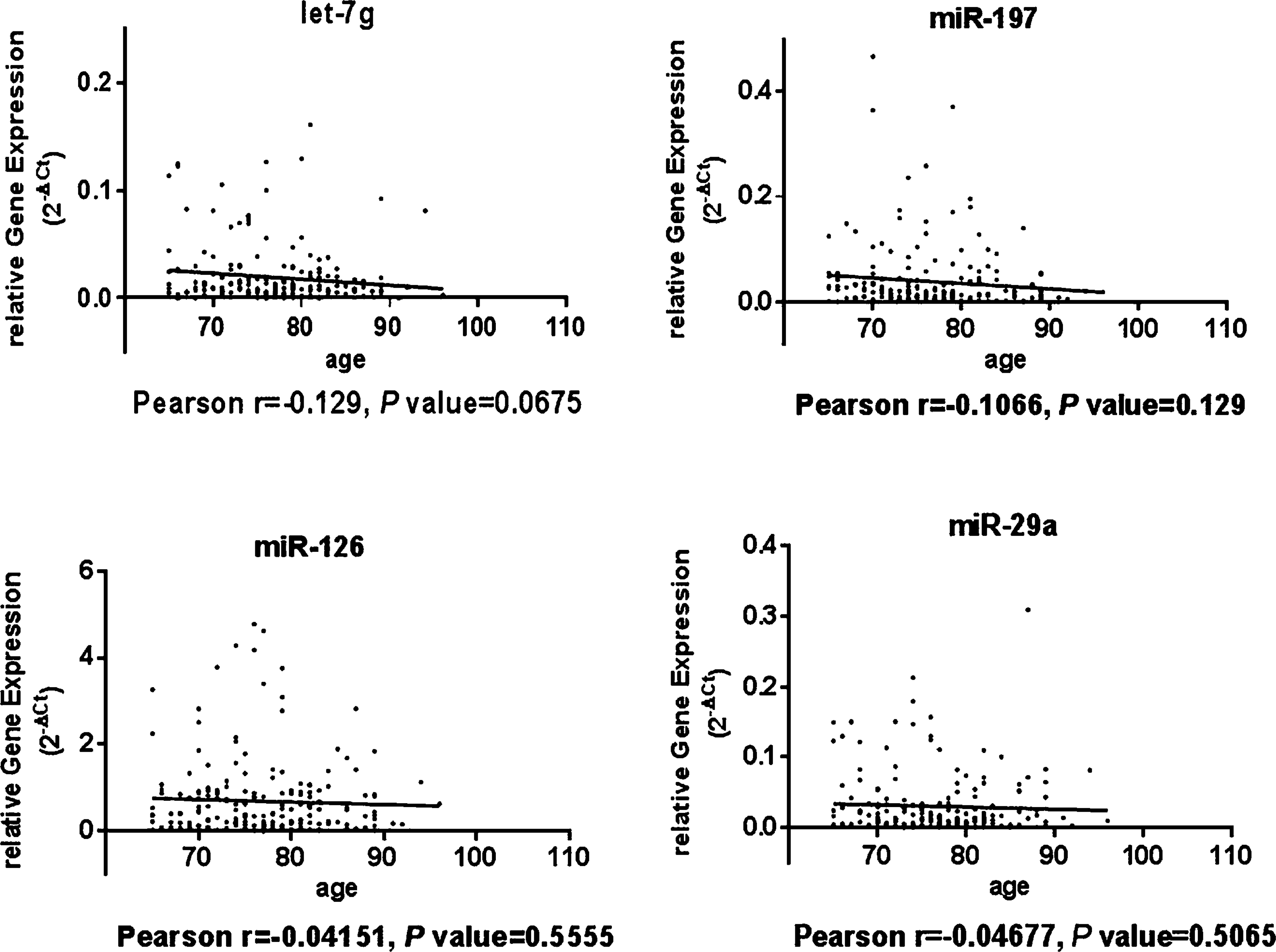
The person correlation between expression of microRNAs and age. Pearson correlation analysis showed let-7g, miR-197, miR-126 and miR-29a had no significant correlations with age (*P*>0.05).

**Figure 4:**
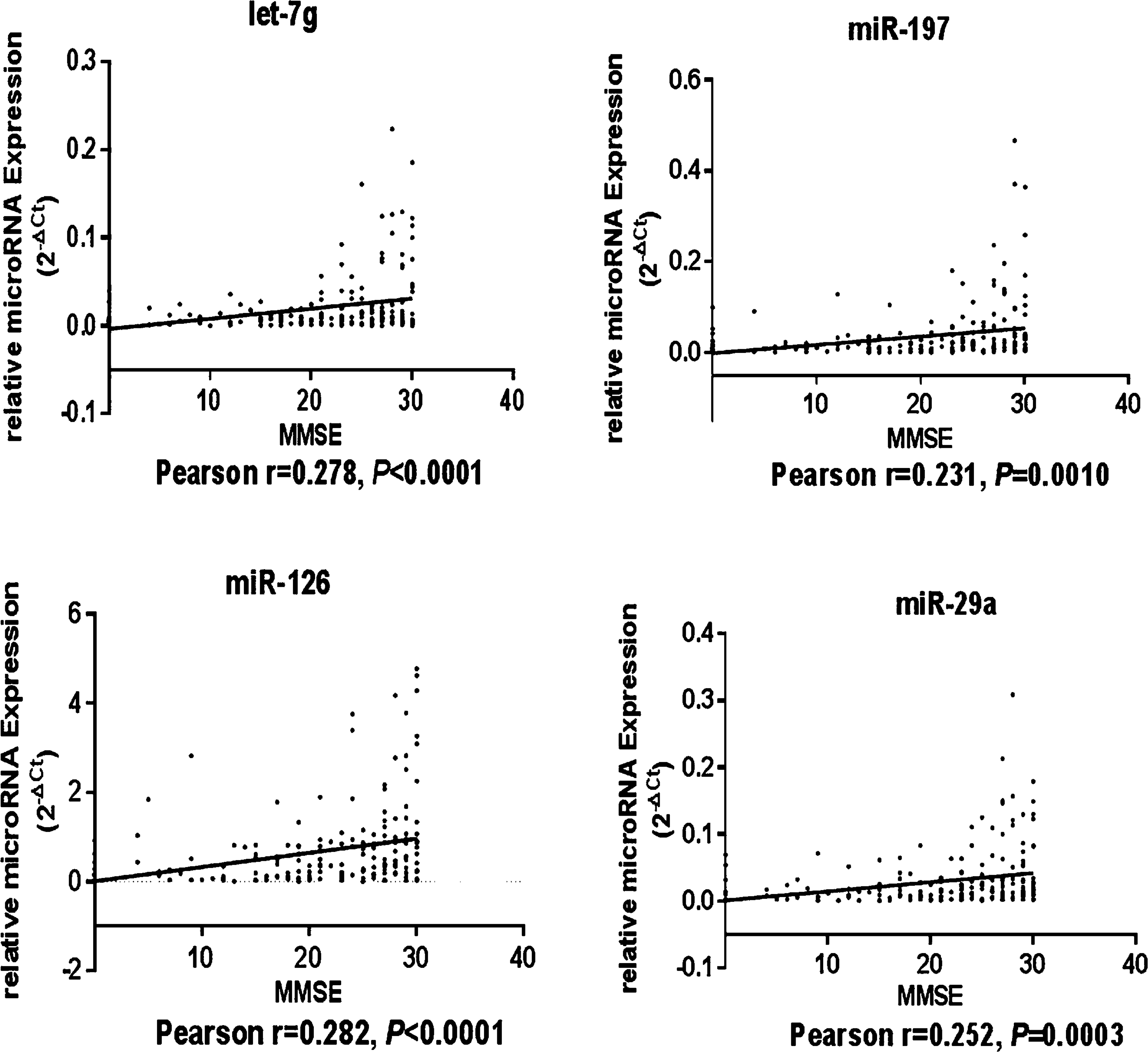
The person correlation analysis among the expressions of let-7g, miR-197, miR-126 and miR-29a and MMSE score. Pearson correlation analysis showed let-7g, miR-197, miR-126 and miR-29a had significant positive correlations with MMSE score (Pearson r=0.29, 0.25, 0.28 and0.22 respectively, *P*<0.001).

### Establishing the predictive microRNA Panel

A stepwise logistic regression model was applied on the training data set (150 participants, 50 participants from each group) to estimate the risk of being diagnosed with MCI and AD. The diagnostic microRNA panels were established based on the logistic regression model for differentiating MCI and AD from CNC. The logit model was constructed based on the four microRNA panel (let-7g, miR-197, miR-126 and miR-29a). The predicted probability of being diagnosed with MCI and AD from the _logit model, Logit (Y=MCI)=*e*_1_·_52 – 29_·_45 × let-7g – 5_·_26 × miR-197 – 0_·_55 × miR-126 -14_·_04 × miR-29a_/(1+ *e*_1_·_52 – 29_·_45 × let-7g – 5_·_26 × miR-197 – 0_·_55 × miR-126 -14_·_04 × miR-29a_) and Logit (Y=AD)=*e*_1_·_49 – 20_·_92 × let-7g – 12_·_62 × miR-197 – 0_·_65 × miR-126 – 6_·_0 × miR-29a_/(1+ *e*_1_·_49 – 20_·_92 × let-7g^−12^·^62 × miR-197 – 0^·^65 × miR-126 – 6^·^0 × miR-^29^a^) were used to construct the receiver operating characteristic (ROC) curve. The diagnostic performance for the two established microRNA panels was evaluated by using ROC analysis. Area under the ROC curve (AUC) was used as an accuracy index for evaluating the diagnostic performance of the selected microRNA panel. The AUC of the microRNA panel was 0.79 (95% CI,0.70 to 0.88) with 84.0% sensitivity and 60.0% specificity in differentiating MCI from CNC (Figure 5A) and the AUC of the microRNA panel on diagnosing AD from CNC was 0.77 (95% CI, 0.68 to 0.86) with 92.0% sensitivity and 50.0% specificity (Figure 5B). Then the microRNA panel (let-7g, miR-197, miR-126, miR-29a) combined with MMSE score was established, logit (Y=MCI)=*e*^17^·^43 – 49^·^09 × let-^7^g –^0^·^30^× miR-197 –1^·^20 × miR-^ 126 - 2_·_08 × miR-29a – 0_·_61 × MMSE _/(1+ *e*_17_·_43 – 49_·_09 × let-7g –0_·_30× miR-197 –1_·_20 × miR-126 - 2_·_08 × miR-29a –0_·_61 × MMSE_) and (Y=AD)=*e*_24_·_28 – 2_·_64 × let-7g – 18_·_41 × miR-197 – 0_·_49 × miR-126 – 8_·_05 × miR-29a0.95× MMSE_/(1+ *e*_24_·_28 – 2_·_64 × let-7g – 18_·_41 × miR-197 – 0_·_49 × miR-126 – 8_·_05 × miR-29a – 0_·_95 × MMSE) wereused to construct the ROC curve. The diagnostic performance of the microRNA panel combined with MMSE score was better than that of the microRNA panels without MMSE score evaluated by using ROC analysis. The AUC for the microRNA panel combined with MMSE score was elevated to 0.93 (95%CI, 0.88 to 0.98) with 75.5% sensitivity and 97.9% specificity in differentiating MCI from CNC (Figure 5C) and the AUC for the microRNA panel combined with MMSE score on diagnosing AD from CNC was increased to 0.99 (95% CI, 0.96 to 1.00) with 97.9% sensitivity and 97.9% specificity (Figure 5D).

**Figure 5:**
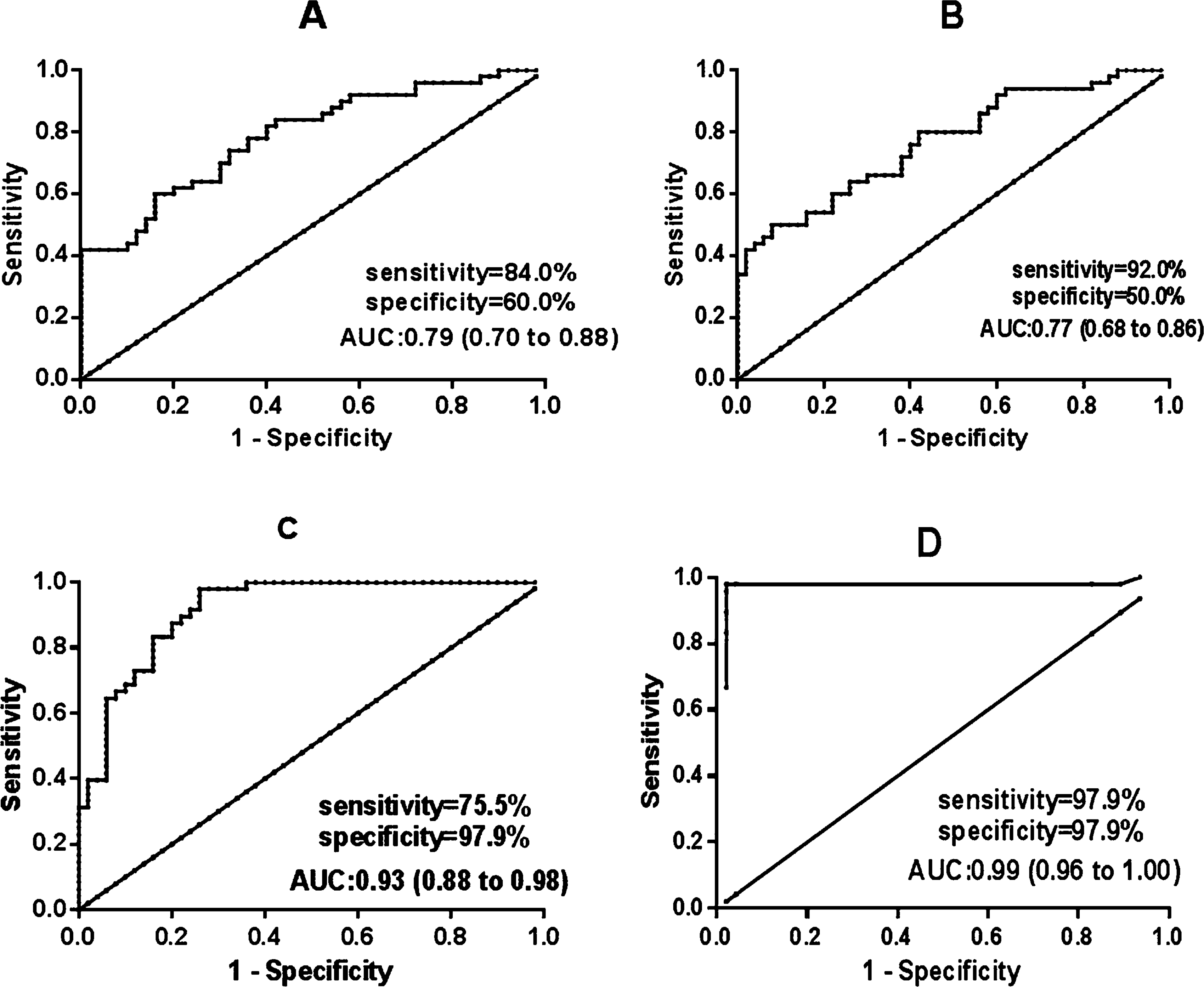
Receiver operating characteristic (ROC) curve analysis of microRNA panel and microRNA panel combined with MMSE in differentiating MCI and AD from CNC in the training set. (A) Area under the ROC curve (AUC) estimation for the microRNA panel in differentiating MCI from CNC. The AUC of the microRNA panel was 0.79 (95% CI,0.70 to 0.88) and the sensitivity and specificity was 84.0% and 60.0%, respectively. (B) AUC estimation for the microRNA panel in differentiating AD from CNC. The AUC of the microRNA panel was 0.77 (95% CI, 0.68 to 0.86) and the sensitivity and specificity was 92.0% and 50.0%, respectively. (C) AUC estimation for the microRNA panel combined with MMSE in differentiating MCI from CNC. The AUC of the microRNA panel was 0.93 (95% CI, 0.88 to 0.98) and the sensitivity and specificity was 75.5% and 97.9%, respectively. AUC estimation for the microRNA panel combined with MMSE in differentiating AD from CNC. The AUC of the microRNA panel was 0.99 (95% CI, 0.96 to 1.00) and the sensitivity and specificity was 97.9% and 97.9%, respectively. Values represent mean ± SEM. CNC, n=50; MCI, n=50; AD, n=50.

### Validating the microRNA panel

The parameters estimated from the training data set were used to predict the probability of being clinically diagnosed with MCI and AD from CNC for the independent validation data set (fifty two participants, eighteen participants from CNC and MCI group, sixteen participants from AD group). The predicted probability was used to construct the ROC curve. The AUC of the four microRNA panel (let-7g, miR-197, miR-126 and miR-29a) was 0.82 (95%CI, 0.68 to 0.96) with 88.9%sensitivity and 66.7% specificity in differentiating between MCI and CNC (Figure 6A) and the AUC of the microRNA panel on diagnosing AD from CNC was 0.82 (95%CI,0.69 to 0.96) with 81.3% sensitivity and 66.7% specificity (Figure 6B). When combined with MMSE score, the AUC of the microRNA panel in differentiating between MCI and AD from CNC were was further improved to 0.91 and 1.00 (Figure 6C and 6D).

**Figure 6:**
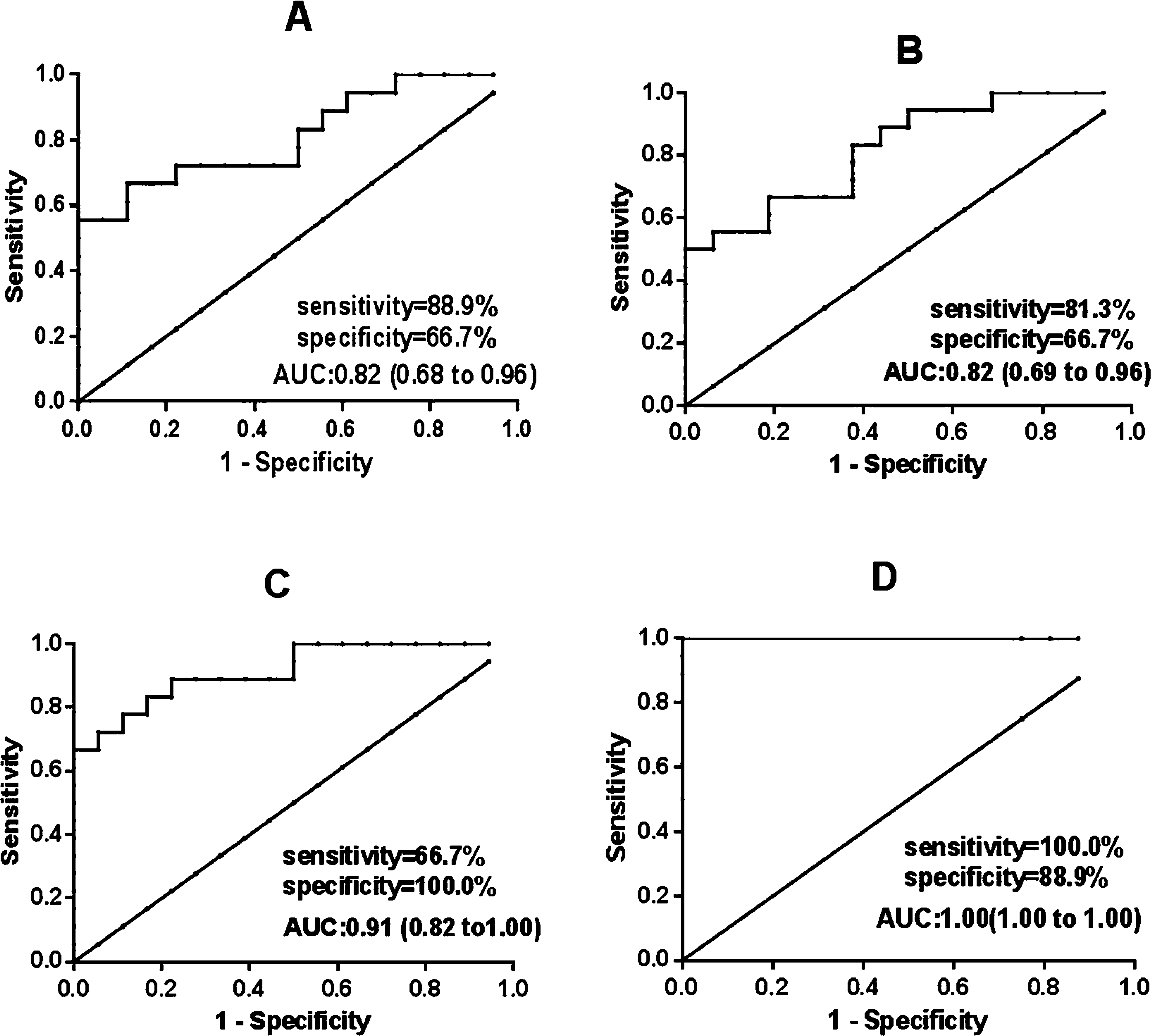
ROC curve analysis of microRNA panel and microRNA panel combined with MMSE in differentiating MCI and AD from CNC in the validation set. (A) AUC estimation for the microRNA panel in differentiating MCI from CNC. The AUC of the microRNA panel was 0.82 (95% CI, 0.68 to 0.96) and the sensitivity and specificity was 88.9% and 66.79%, respectively. (B) AUC estimation for the microRNA panel in differentiating AD from CNC. The AUC of the microRNA panel was 0.82 (95% CI, 0.69 to 0.96) and the sensitivity and specificity was 81.3% and 66.7%, respectively. (C) AUC estimation for the microRNA panel combinedwithMMSEin differentiating MCI from CNC. The AUC for the microRNA panel was 0.91 (95% CI, 0.82 to 1.00) and the sensitivity and specificity was 66.7% and 100.0%, respectively. (D) AUC estimation for the microRNA panel combined with MMSE in differentiating AD from CNC. The AUC for the microRNA panel was 1.00 (95% CI, 1.00 to 1.00) and the sensitivity and specificity was 100.0% and 88.9%, respectively. Values represent mean ± SEM. CNC, n=18; MCI, n=18; AD, n=16.

### MicroRNAs target genes prediction and Pathway enrichment

To determine possible target genes for the differentially expressed microRNAs, we searched the TargetScan 6.2 (http://www.targetscan.org) database and Pictar (http://pictar.bio.nyu.edu/) for predicted microRNA targets in mammals. A total of 2689 genes were predicted as targets for the four microRNAs from TargetScan. The target genes were 1180, 1201, 25 and 283 for let-7g, miR-29a, miR-126 and miR-197, respectively. While Pictar predicted total 660 target genes (310, 186, 134 and 30 target genes for let-7g, miR-29a, miR-126 and miR-197). Two hundred and thirty one microRNA target genes were cross validated from both TargetScan and Pictar (Supplement Table 4). Thus, the results suggested that the human microRNA target genes regulated by an individual microRNA generally constitutes the biological network of functionally-associated molecules (Satoh & Tabunoki, 2011). Therefore, it is possible that even small changes in the expression of a single microRNA could affect a wide range of signaling pathways and networks involved in diverse biological functions. To identify biologically relevant molecular networks from the target genes,we could analyze them by using pathway analysis tool of bioinformatics endowed with comprehensive knowledgebase, such as KEGG. KEGG includes manually curated reference pathways that cover a wide range of metabolic, genetic, environmental, and cellular processes, and human diseases. Thus, these target genes were submitted to KEGG database for pathway illustration. The target genes constructed KEGG pathways related to various types of cancers (small cell lung cancer, glioma, melanoma and Pancreatic cancer), focal adhesion, and signaling pathways of p53, MAPK and insulin (Supplement Table 5). These results suggested an existence of collaborative regulation of gene expression by transcription factors and microRNAs in cancer-associated microRNA target gene networks.

## Discussion

Though several methods have been developed to alleviate the symptoms of AD, no therapies have been developed that can modify the course of the disease (Huang & Mucke, 2012). This stems not only from a lack of a detailed understanding of the pathophysiological processes underpinning the disease, but also an inability to diagnose patients both accurately and early. It has been estimated that about two-thirds of dementia patients go undiagnosed (Ho et al, 2010), and that by the time of diagnosis AD pathology has been developing for˜20 years (Villemagne et al, 2013). Thus biomarkers that can diagnose AD accurately and early are urgently needed to improve the diagnostic sensitivity and specificity and to monitor the biological activity of AD in terms of the burden of neuronal involvement and the rate of disease progression.

Current methods for the diagnosis of AD depend on a combination of brain imaging and tests of cerebrospinal fluid (CSF) and blood to identify changes that signal AD. However, the imaging techniques do not afford sufficient accuracy in discriminating between AD and non-AD, especially between MCI and non-AD dementia to provide clear and decisive diagnosis (Morris et al, 2016) and wide variations in sensitivity and specificity have been reported. The diagnostic use of CSF is limited because of invasive collection and the potential side effects by lumbar puncture. Thus, the ability to identify biomarkers through less invasive procedures, such as in a blood test, would be significant. Circulating serum microRNA might represent a novel putative, easily accessible and stable marker. The recent discovery of aberrant expression of microRNAs in AD provided clues for analyzing circulating microRNAs for AD diagnosis.

In this study, we revealed that the expression levels of let-7g, miR-197, miR-126 and miR-29a in serum were decreased significantly in MCI and AD group than those of CNC group. Furthermore, the expression levels of the four microRNAs were correlated with MMSE score, which suggested that they were potential circulating biomarkers for diagnosing MCI and AD. The microRNA panel with the four microRNAs from the multivariate logistic regression model demonstrated good diagnostic performance in differentiating MCI and AD from CNC. When combined with MMSE score, the diagnostic performance of the microRNA panel was further improved.

Among the microRNA panel with four microRNAs observed in our study, only miR-29 has been previously reported the association with AD. Several groups have revealed changes in miR-29 expression levels in the AD brain, where a decrease in miR-29 was observed in most of the studies (Lugli et al, 2015; Shioya M, 2010). This decrease was also observed in the cerebral cortex of transgenic mice bearing APP and PS1 familial AD mutations (Wang et al, 2009). Interestingly, miR-29 has been shown to target BACE1 which is directly implicated in AD pathogenesis through the formation of toxic Aβ peptides in vitro and in vivo (Hebert et al, 2008; Lei et al, 2015). These observations suggest a potential role for miR-29 in the pathogenesis of AD. Besides miR-29, the other three microRNAs (let-7g, miR-197, and miR-126) have not been identified or investigated in relationship to AD till now. Only one research reported that miR-197 was downregulated in white matter, let-7g and miR-126 were downregulated in gray matter of AD (Zong et al, 2011). Moreover, miR-197 was also found differentially expressed in cerebral spinal fluid of AD patients (Cogswell et al, 2008). MiR-126 is highly enriched in endothelial cells and plays a pivotal role in maintaining endothelial homeostasis and vascular integrity (Wang et al, 2008), regulating expression of adhesion molecules and controlling vascular inflammation (Harris et al, 2008). High expression of miR-126 has been reported in the human and rodent brain (Landgraf et al, 2007) and in cultured rat motoneurons (Wei et al, 2010) indicated that it could play a role in neuronal cell function. Potential role of miR-126 in the nervous disorders could be related with its profound impact on regulating factors in key signaling pathways that are associated with disease processes. For example, miR-126 is involved in insulin-like growth factor (IGF)-1, vascular epithelial growth factor or EGF-like domain containing protein 7 (EGFL7) signaling. Insulin/IGF-1 signaling promotes neurite sprouting, synapse formation, and neuronal survival, and it has been implicated in memory, aging, and neurodegeneration (Bishop et al, 2010). In AD, neurons are resistant to IGF-1 and there is a large body of data linking changes in growth factor/ IGF-1 signaling to disease pathogenesis and altered neuronal cell function (Freude et al, 2009). Another way that miR-126 functions in the nervous system could be through its genomic and regulatory link with EGFL7 (Sonntag et al, 2012). MiR-126 is encoded in intron 7 of the EGFL7 gene and originates from the EGFL7 pre-mRNA (Wang et al, 2008). EGFL7 is expressed in the mammalian brain where it regulates notch signaling in neural stem cells altering their self-renewal and multipotency as well as their differentiation potential toward neuro- and gliogenesis (Schmidt et al, 2009). EGFL7 could have implication for boosting adult neurogenesis as a repair mechanism in neurodegenerative disorders and brain injury (Bicker & Schmidt, 2010). Although it remains to be determined whether miR-126 is involved in these processes, given its impact on IGF-1 and EGFL7 regulation in a magnitude of cell systems, it is likely that similar mechanisms also occur in neurons.

Because a single microRNA concurrently downregulates hundreds of target mRNAs, the set of microRNA target genes coregulated by an individual microRNA generally constitutes the biologically integrated network of functionally associated molecules(Hsu et al, 2008; Satoh & Tabunoki, 2011). Even small changes in the expression level of a single microRNA could affect a wide range of signaling pathways involved in diverse biological functions. From this point of view, the characterization of a global picture of microRNA target networks would promote us to understand microRNA-mediated molecular mechanisms underlying AD. To identify biologically relevant molecular networks and pathways of great magnanimity data, pathway analysis tool of bioinformatics with comprehensive knowledgebase, such as KEGG analyze could be used. KEGG is a public database, including manually curated reference pathways that cover a wide range of metabolic, genetic, environmental, and cellular processes, and human diseases. Currently, KEGG contains 198,560 pathways generated from 428 reference pathways. From the results of target gene-pathway enrichment, we found that the target genes constructed KEGG pathways were related to various types of cancers, such as small cell lung cancer, glioma, melanoma and pancreatic cancer. Although microRNA has been actively researched in cancer for a number of years, this is a relatively new area for research in AD and other neurodegenerative diseases. In recent years, more and more researches show that many microRNAs have differential expressions in both AD and a variety of cancers (KC, 2010; Wu et al, 2008). This phenomenon was also observed in our results. Mir-197 and let-7g are involved in cancer initiation and progression, which were identified by several studies (Hamada et al, 2013; Kumar et al, 2008). In our study, their expressions were decreased significantly in MCI and AD group than those of CNC group. The aberrant expressions of mir-197 and let-7g in human AD brain indicated that they might play crucial roles in AD as well as cancer (Hebert et al, 2008; Zong et al, 2011). The increasing evidences for overlapping microRNA functions in cancer and neurodegeneration(Du & Pertsemlidis, 2011) suggest that microRNAs play multiple regulatory roles in pathways active across both diseases and the molecular machinery involved in maintaining neural function in neurodegenerative disease may be shared with oncogenic pathways (Saito & Saito, 2012). MicroRNAs appear to be important in the common signaling pathways that regulate differentiation, proliferation and death of cells in both cancer and AD and may function using some of the same mechanisms(Holohan et al, 2012). Some results indicate that chromatin remodeling by epigenetic treatment can directly modulate expression of microRNAs that are involved in the pathogenesis of cancer and neurodegenerative disease. Nevertheless, there are very little functional data of mir-197 and let-7g in the central nervous system, much work remains to be accomplished to elucidate the roles of them in both normal and AD status.

To date, it is becoming clear that the consideration of a single biomarker might not be potent enough to improve diagnostic specificity. It seems probable that only the combination of several biomarkers derived from blood will be successful to define a patient-specific signature (Humpel, 2011). Thus, circulating microRNAs, especially microRNA panel may be used as a potential novel noninvasive biomarker for AD diagnosis and prognosis assessment. Although there is a long history of investigation of circulating mRNA molecules as potential biomarkers, blood-based microRNA studies are in their infancy. In this study, we revealed differentially expressed microRNAs in serum from MCI, AD patients and healthy controls in a large population-based cohort and construct microRNA panel that demonstrated good diagnostic performance for both MCI and AD. When combined with MMSE score, the diagnostic performance of the microRNA panel was further improved, which suggested that microRNA profile along with other biomarkers and cognitive tests could potentially provide a comprehensive and non-invasive methodology for the early diagnosis of AD. The result that the target genes pathways relating to various types of cancers demonstrated possible intersections between microRNA functions in cancer and AD, suggesting that a similar strategy could be useful in studying both AD and cancer. The microRNAs in cancer and AD could benefit from future research in both diseases. Although our study got encouraging results of serum microRNA panel in early diagnosis of AD, it must be noted that the blood serum samples used in thisstudy were only clinically diagnosed and further studies are necessary to assess their ability to discriminate them from other forms of dementia. In addition, independent validation of these microRNAs as biomarkers will also be required. Whether these microRNA profiles could provide conclusive diagnosis is currently unknown, nonetheless, microRNA profiles along with other biomarkers and cognitive tests could potentially provide a more comprehensive and early diagnosis and prognosis of AD. Therefore, further studies and additional work will be needed to assess the ability and applicability in the future.

## Materials and methods

### Patients

The 60 serum samples (including 20 CNC, 20 MCI and 20 AD) in discovery phase and 202 serum samples (including 68 CNC, 68 MCI and 66 AD) in training and validation phases that met the eligibility criteria, were collected from neurology department of Chinese People’s Liberation Army (PLA) General Hospital and Beijing Geriatric Hospital, recruited between May 2008 and March 2011. The samples have been clinically diagnosed by neurologists, neuropsychologists, and other staff members in the neurology department of Chinese PLA general hospital and Beijing Geriatric Hospital according to NINCDS/ADRDA criteria (McKhann G, 1984) and MCI according to the Petersen criteria (Petersen et al, 1999).

### Study Design

This study included three phases. In each study phase, blood samples were obtained from participants including CNC, MCI and AD patients. Studies were approved by the Ethics Committee of Chinese PLA general Hospital and Beijing Geriatric Hospital. Informed consent was obtained from all participants.

#### 1. Discovery phase

Sixty serum samples from CNC, MCI and AD patients, each with 877 microRNAs, were screened with a microarray platform. The patient characteristics were presented in Table 1. A Mann-Whitney test was performed to discover differentially expressed microRNAs in the three pairwise comparisons: CNC versus AD and MCI, respectively.

From the 8 differentially expressed microRNAs, fourteen detectable microRNAs with fold change > 2.6 and *P* value < 0.037 were tested by qRT-PCR, with cohort of thirty participants.

#### 2. Training phase

The 14 microRNAs discovered via microarray were first tested with qRT-PCR in an independent cohort of serum samples from 30 participants. Five microRNAs (let-7g, miR-197, miR-126, miR-29a and miR-451) were detected in all groups and passed the quality control. The expressions of miR-451 were not different in CNC, MCI and AD groups. Thus, four microRNAs (let-7g, miR-197, miR-126 and miR-29a) that were differentially expressed among AD, MCI and CNC groups were further tested in an additional 120 participants. Finally, the 150 participants were used as the training set to construct the diagnostic microRNA panel based on the logistic regression model for the differentiation not only between the AD group and the CNC group but also between the MCI group and the CNC group. The patient characteristics in the training phase were presented in Table 1.

#### 3. Validation phase

The parameters of the logistic model from the training phase were applied to an independent cohort of fifty two samples for validating the diagnostic performance of the selected microRNA panel. The patient characteristics in the validation phase were presented in Table 1.

### Blood serum collection and RNA Isolation

Whole blood was centrifugation at 1600 g for 15 min following 30 min of clotting at room temperature. Serum samples were transferred into a new tube and stored at - 80 ^°^C. Total RNA was extracted by using mirVana PARIS microRNA Isolation kit according to the instructions from the manufacturer (Ambion, Austin, TX). The concentration was quantified by NanoDrop 2000c Spectrophotometer (NanoDrop Technologies, Waltham, MA).

### RNA isolation

Total RNA was extracted from the serum by using mirVana PARIS microRNA Isolation kit according to the instructions from the manufacturer (Ambion, Austin, TX). The concentration was quantified by NanoDrop 1000 Spectrophotometer (NanoDrop Technologies, Waltham, MA).

### Genome-wide microRNA analysis

Human microRNA microarrays (Agilent Technologies, Santa Clara, CA) were used in 60 serum samples (n=20 normal, n=20 MCI and n=20 AD) to compare the expression profiles. The microarray contains probes for 877 human microRNAs from Sanger database v.14.0. Each slide is formatted with eight identical arrays. Total RNA (100 ng) derived from plasma samples were labeled with Cy3. Microarray slides were scanned by XDR Scan (PMT100, PMT5). The labeling and hybridization were performed according to the protocols in the Agilent microRNA microarray system. The microarray image information was converted into spot intensity values using Feature Extraction Software Rev. 9.5.3 (Agilent Technologies, Santa Clara, CA). The signal after background subtraction was exported directly into the GeneSpring GX10software (Agilent Technologies, Santa Clara, CA). The raw signals obtained for single-color CY3 hybridization were normalized by a stable endogenous control miR-1228. After that, a log transform with base 2 was performed. A sample that showed intra-array coefficients of variation (CV) across replicated spots on an array above 15% or positive signals less than 5% was considered to be unreliable and excluded from further analysis. A detectable microRNA was defined as the microRNA had positive signals on microarrays in > 50% of plasma samples from any one of the four category subjects.

### Quantitative RT-PCR

For testing of candidate microRNAs acquired on microarrays, qRT-PCR was performed using Taqman microRNA assays (Applied Biosystems, Foster City, CA).

Assays to quantify mature microRNAs were conducted using a Taqman microRNA PCR kit (Applied Biosystems, Foster City, CA, USA) according to the manufacturer’s instructions. Briefly, 8 ng total RNA was reverse-transcribed to cDNA and looped anti-sense primers (Applied Biosystems, Foster City, CA, USA). The mixture wasincubated at 16 ^°^C for 30 min, 4 ^°^C for 30 min and 85 ^°^C for 5 min to generate microRNA cDNAs. Real-time PCR was subsequently performed using an Applied Biosystems 7500 Sequence Detection System (Applied Biosystems, Foster City, CA,USA). The reactions were incubated in a 96-well optical plate at 95 ^°^C for 10 min followed by 40 cycles consisting of a 15 s interval at 95^°^C and a 1 min interval at 60 ^°^C. The expression level of miR-16 was used as a stable endogenous control for normalization. All assays were carried out in triplicate.

### MicroRNA target gene prediction

The TargetScan algorithm was used to identify predicted gene targets of the differentially expressed microRNAs. TargetScan was mainly selected because it predicted the highest number of known microRNA targets obtained from TarBase 5.0 (http://microrna.gr/tarbase) in our subset of differentially expressed microRNAs (Papadopoulos et al, 2009).

Potential target genes of microRNA were analyzed by the currently available major prediction algorithms, including TargetScan (http://targetscan.org) and PicTar (http://pictar.bio.nyu.edu).

### KEGG pathway

Pathway analyses of the differentially expressed genes were based on the Kyoto Encyclopedia of Genes and Genomes (KEGG). KEGG pathway database is a recognized and comprehensive database including all kinds of biochemistry pathways(Kanehisa & Goto, 2000). In this work, the KEGG database was applied to investigate the enrichment analysis of the differentially expressed genes to find the biochemistry pathways which might be involved in the occurrence and development of AD.

### Statistical analysis

For microarray analysis, the Mann-Whitney unpaired test was used for the three pairwise comparisons (AD *vs* CNC, MCI *vs* CNC, and MCI *vs* AD). For the data obtained by qRT-PCR, the Mann-Whitney unpaired test was used for the comparison among the three groups. A stepwise logistic regression model was used to select diagnostic microRNA markers based on the training dataset. The predicted probability of being diagnosed with MCI and AD were used as a surrogate marker to construct receiver operating characteristic (ROC) curve. Area under the ROC curve (AUC) was used as an accuracy index for evaluating the diagnostic performance of the selected microRNA panel. SAS (version 10.4.7.0) software and GraphPad prism (version 5.0) were used to perform the ROC and regression analysis. All *P* values were two sided.

## Acknowledgments

We thank Professor Ying Wu (Shanghai Medical College, Fudan University) for the support on data analysis and interpretation and our colleagues (Si-Di Li, Tao Wang, Yi-Zhou Mei, Feng Liu and Yan Sun) for their work on recruiting patients and collecting serum samples.

## Author contribution

Ning Jiang was second applicant for the grant of The National High technology Research and Development Program of China (2008AA02Z423), did the repackaging clinical samples, the microRNA screening by the Human microRNA microarrays 14.0 of this study, study design, data collection, data interpretation, writing and revising the manuscript.

Can-Jun Ruan did the literature search, all experiments, study design, data collection, data statistical analysis, data interpretation, and writing the manuscript.

Xiao-Rui Cheng did data collection, data statistical analysis and data interpretation.

Lu-Ning Wang was principal investigator for the clinical cases recruitment and serum samples collection.

Ji-Ping Tan and Wei-Shan Wang recruited patients, and were responsible for collecting serum samples.

Fang Liu did the repackaging clinical samples and microRNA data analysis.

Wen-Xia Zhou was lead applicant for grants of National Science and Technology Major Project (2012ZX09301003-002). Yong-Xiang Zhang was lead applicant for the grant of The National High technology Research and Development Program of China (2008AA02Z423). Wen-Xia Zhou and Yong-Xiang Zhang were principal investigators for this study, and responsible for last draft of the manuscript.

## The Paper Explained

PROBLEM: Early, reliable and non-invasive diagnosis of Alzheimer disease (AD) is a great challenge, which makes the prognosis and therapeutic interventions quite difficult. Blood microRNAs are promising biomarkers that could aid early diagnosis and ultimately lead to the development of more effective interventions. However, most studies are small and relatively heterogeneous. Mild cognitive impairment (MCI) is defined as the intermediary state between normal old age cognition and AD which reflects preclinical AD in a considerable portion of the affected individuals. Therefore, in this study, we aimed to investigate the expressions of microRNA in serum and identify microRNA panel for diagnosing MCI and AD.

RESULTS: Serum microRNA expression was investigated with two independent cohorts including 202 participants non-dementia control, MCI, and AD. Quantitative reverse-transcriptase polymerase chain reaction assay was applied to evaluate the expression of four microRNAs (let-7g, miR-197, miR-126 and miR-29a) which were screened from 877 microRNAs in 60 serum samples with a microarray platform in prior study. Logistic regression model based on microRNA panel was constructed using a training cohort (n=150) and then validated using an independent cohort (n=52). Area under the receiver operating characteristic curve was used to evaluate diagnostic accuracy. The expression levels of four microRNAs were significantly decreased in MCI and AD groups, and positively correlated with mini mental state examination (MMSE) score. Then, we identified a microRNA panel (let-7g, miR-197, miR-126 and miR-29a) that demonstrated good diagnostic performance for MCI and AD. When combined with MMSE score, the diagnostic performance of the microRNA panel was further improved.

IMPACT: We found a serum microRNA panel that has considerable clinical value in diagnosing AD. Four microRNAs were potential circulating biomarkers in diagnosing AD.

